# Fetal therapy model of myelomeningocele with three-dimensional skin using amniotic fluid cell-derived induced pluripotent stem cells

**DOI:** 10.1101/095281

**Authors:** Kazuhiro Kajiwara, Tomohiro Tanemoto, Seiji Wada, Jurii Karibe, Norimasa Ihara, Yu Ikemoto, Tomoyuki Kawasaki, Yoshie Ohishi, Osamu Samura, Kohji Okamura, Shuji Takada, Hidenori Akutsu, Haruhiko Sago, Aikou Okamoto, Akihiro Umezawa

## Abstract

Myelomeningocele (MMC) is a congenital disease without genetic abnormalities. Neurological symptoms are irreversibly impaired after birth. No effective treatment has been reported to date. Only surgical repairs have reported so far. In this study, we performed antenatal treatment of MMC with an artificial skin using induced pluripotent stem cells (iPSCs) generated from a patient with Down syndrome (AF-T21-iPSCs) and twin–twin transfusion syndrome (AF-TTTS-iPSCs) to a rat model. We manufactured three-dimensional skin with epidermis generated from keratinocytes derived from AF-T21-iPSCs and AF-TTTS-iPSCs and dermis of human fibroblasts and collagen type I. For generation of epidermis, we developed a novel protocol using Y-27632 and epidermal growth factor. The artificial skin was successfully covered over MMC defect sites during pregnancy, implying a possible antenatal surgical treatment with iPSC technology.

## Introduction

Myelomeningocele (MMC) is the most common neural tube defect characterized by a skin defect in addition to defects of the midline vertebra and dura mater. Under these conditions, the spinal cord is exposed to the external environment. Although MMC is compatible with survival, this condition ranks as one of the most severe birth defects, with the manifestation of sequelae that affect both the central and peripheral nervous systems, leading to lifelong paralysis and dysfunction of the bladder and bowel for which no cures exist. The two-hit hypothesis postulates irreversible neurological deficits during pregnancy. First, a failure of primary neurulation results in the neural tube defect. Second, the subsequent destruction of the exposed neural tissue is attributed to the chemical damage by amniotic fluid (AF) containing meconium and to the physical damage caused by the direct trauma (Heffez et al., 1990; McLone and Knepper, 1989; Michejda, 1984). Therefore, fetal intervention is focused to prevent progressive neurological damage *in utero* (Bruner et al., 2004; Johnson et al., 2003; Sutton et al., 1999; Tubbs et al., 2003; Tulipan et al., 1999; Tulipan et al., 2003; Walsh et al., 2001). The first human randomized-controlled clinical trial entitled the “Management of Myelomeningocele Study (MOMS)” has found that intrauterine repair of fetal MMC improves the neurological prognosis compared with postnatal MMC repair (Adzick et al., 2011). However, the MOMS trial also proved that severe complications such as preterm birth, premature rupture of the membrane, and placental abruption and uterine wall dehiscence at the repair site increased due to a large incision of the uterine wall. Moreover, the skin defect is too large to allow skin closure in 20%–30% of fetuses with MMC (Mangels et al., 2000). Although several less invasive methods that prevent leaking of cerebral spinal fluid with coverage of the exposed spinal cord using various coverage materials, such as gelatin sponge incorporating basic fibroblast growth factor and autologous amniotic membrane, have been reported, improvement of neurological function remains insufficient because insufficient recovery of neural damage occurs during pretreatment (Brown et al., 2014; Kohl et al., 2006; Kohl et al., 2009; Watanabe et al., 2010; Watanabe et al., 2011). Thus, the development of a less invasive method is required to be applied during earlier gestation.

Coverage of a skin defect by a three-dimensional (3D) skin enables fetal intervention to be not only less invasive but also completed by an earlier term of pregnancy because autologous cultured skin transplantation enhance wound healing such as autologous skin transplantation for patients with severe burns (Pham et al., 2007). Moreover, 3D skin may grow in harmony with the fetal growth during all pregnancy periods. However, it is difficult to provide available autologous skin grafts for a fetus with MMC. Furthermore, the strong demand far outstrips the current supply of skin graft. Therefore, we hypothesize that induced pluripotent stem cells (iPSCs) can be an ideal new material for autologous skin transplantation. The first clinical therapy using human iPSCs treated a Japanese woman with macular degeneration (Sivan et al., 2016). A great deal of research is currently in progress to facilitate therapy for patients with intractable diseases, such as Parkinson’ disease and spinal cord injury (Cherry and Daley, 2012, 2013). iPSCs can provide therapeutic potential for tissue repair and regeneration in combination with *in vitro* differentiation. The use of iPSCs in clinical application has largely been welcomed by society because the use of these cells avoids the substantial ethical concerns and immune rejection.

In the present study, we generated human iPSCs from amniotic fluid cells (AFCs) using feeder-free, xeno-free, and integration-free systems. To verify that iPSCs could be generated from fetuses with severe fetal disease, we chose two major fetal diseases, twin–twin transfusion syndrome (TTTS) and trisomy 21 (T21): TTTS is one of the most serious complications of monochorionic multiple gestations; fetal intervention is most frequently performed, and trisomy 21 is the most common chromosome abnormality among liveborn infants. To apply iPSC-based cell therapy during earlier gestational age, we established an effective protocol for differentiation of iPSCs into keratinocytes by the addition of Y-27632 and epidermal growth factor (EGF). We successfully established a novel surgical approach in the fetal rat model of MMC using reconstructed 3D skin with iPSC-derived keratinocytes (iPSC-KC) and investigated the effect of transplantation of 3D skin.

## Materials and Methods

### Ethical statement

The protocol for the use of human cells in the present study was approved by the Institutional Review Board of the National Research Institute for Child Health and Development of Japan and was in full compliance with the Ethical Guidelines for Clinical Studies (Ministry of Health, Labor, and Welfare).

### Human cells

AF was obtained from fetus with Down syndrome and TTTS, with both conditions associated with polyhydramnion. In case of TTTS, AF was obtained at the time of fetoscopic laser surgery at gestational ages ranging from the 19th to 26th weeks. In the cases of Down syndrome, AF was obtained by amnioreduction at 29 weeks of gestation. Cells were obtained by the centrifugation of 20-ml AF at 1,500 rpm for 10 min after filtration through a 100-μm filter. The supernatant was removed, and the precipitates were seeded in 60-mm tissue culture dishes, which were precoated with 0.1% gelatin. These cells were incubated at 37°C under 5% humidified CO_2_ in 4 ml of AmnioMAX^TM^-II Complete Medium (Invitrogen, cat. no. 11269-016). Cell clusters were emerged at 6–7 days after seeding. Nonadherent cells were discarded, and the media was changed every 2 days. When the cultures reached subconfluence, the cells were harvested with a trypsin-ethylenediaminetetraacetic acid (EDTA) solution (Wako, cat. no. 209-16941) and replated at a ratio 1:8 in a 60-mm dish. These cells were cultured and passaged routinely at 70%–80% confluence. The cells at passages 3–4 were used for the generation of iPSCs.

HDK1-K4DT, an immortalized keratinocyte, was grown in a keratinocyte serum-free medium (KSFM) (Invitrogen, cat. no. 17005-042) and supplemented with 10 μM Y-27632 (Wako, cat. no. 251-00514) (Egawa et al., 2012). Subconfluent cultures of HDK1-K4DT were passaged at a ratio 1:6. HDK1-K4DT at less than passage 20 was used for organotypic culture. HFF2, an immortalized fibroblast, was grown in Dulbecco’s modified Eagle’s medium (DMEM) supplemented with 10% fetal bovine serum (FBS) (Tatsumi et al., 2006).

### Generation of feeder-free iPSCs

AF-derived cells (AFCs) were seeded at 6 × 10^5^/well in a six-well plate and maintained in AmnioMAX™-II Complete Medium. Three episomal vectors encoding six factors (L-MYC, KLF4, OCT4, SOX2, LIN28, and short-hairpin RNA for P53) (addgene, cat. no. 27077, 27078, 27079) were electroporated into the AFCs on day 0 as previously described (Okita et al., 2011). On day 5, transfected cells were passaged and seeded at 1.3 × 10^6^/dish onto Vitronectin (VTN) (Life Technologies, cat. no. A14701SA)-coated 100-mm plates in Essential 8 (E8) medium (Life Technologies, cat. no. A14666SA), and the medium was changed every 2 days. We observed the appearance of human embryonic stem cell (ESC)-like colonies 30–50 days after electroporation. For feeder-free cultures, plates were coated with 0.5 g/cm^2^ VTN at room temperature for 1 h. iPSCs were maintained in E8 Medium on VTN-coated dishes and passaged using 0.5 mM EDTA in PBS. Colonies with flat, human ESC-like morphology were expanded and maintained successfully for more than 70 passages.

### RT-PCR

Total RNA was isolated from cells using the RNeasy Plus micro Kit (Qiagen, cat. no. 74004), and DNA was removed by DNase treatment (Qiagen, cat. no. 79254). Complementary DNA (cDNA) was synthesized from 1 μg of total RNA using Superscript III reverse transcriptase (Invitrogen, cat. no. 18080-085) with oligo (dT) primer according to the manufacturer’s instructions. Template cDNA was PCR amplified with gene-specific primer sets (Supplemental Table S1).

### Quantitative RT-PCR

RNA was extracted from cells using the RNeasy Plus Micro kit (Qiagen, cat. no. 74004). An aliquot of total RNA was reverse transcribed using an oligo (dT) primer (Invitrogen, cat. no. 18418-20). For the thermal cycle reactions, the cDNA template was amplified (Applied Biosystems^®^ Quantstudio^TM^ 12K Flex Real-Time PCR System) with gene-specific primer sets using the Platinum Quantitative PCR SuperMix-UDG with ROX (Invitrogen, cat. no. 11743-100) under the following reaction conditions: 40 cycles of PCR (95°C for 15 s and 60°C for 1 min) after an initial denaturation (95°C for 2 min). Fluorescence was monitored during every PCR cycle at the annealing step. The authenticity and size of the PCR products were confirmed using a melting curve analysis (using software provided by Applied Biosystems). mRNA levels were normalized using glyceraldehyde-3-phosphate dehydrogenase as a housekeeping gene.

### Immunocytochemical analysis

Cells were fixed with 4% paraformaldehyde (PFA) in PBS for 10 min at 4°C. After washing with PBS and treatment with 0.1% Triton X-100 (Sigma-Aldrich, cat. no. T8787-100ML) for 10 min at 4°C, cells were pre-incubated with 5% normal goat serum (Dako, cat. no. X0907) in PBS for 30 min at room temperature, following which they were reacted with primary antibodies in blocking buffer for 24 h at 4°C. After washing with PBS, cells were incubated with fluorescently coupled secondary antibodies; anti-rabbit or anti-mouse IgG conjugated with Alexa 488 or 546 (1:1000) in blocking buffer for 30 min at room temperature. The nuclei were stained with 4′,6-diamidino-2-phenylindole (DAPI) (Biotium, cat. no. 40043). All images were captured using confocal microscopy (LSM 510 and LSM 510 META laser scanning microscope). Antibody information is provided in Supplemental Table S2.

### Karyotypic analysis

Karyotypic analysis was performed at the Nihon Gene Research Laboratories Inc. (Sendai, Japan). Metaphase spreads were prepared from cells with 100 ng/ml of Colcemid (Karyo Max, Gibco Co. BRL) treatment for 6 h. The cells were fixed with methanol:glacial acetic acid (2:5) three times and dropped onto glass slides (Nihon Gene Research Laboratories Inc.). Chromosome spreads were Giemsa banded and photographed. Twenty metaphase spreads were analyzed for each sample and karyotyped using a chromosome imaging analyzer system (Applied Spectral Imaging, Carlsbad, CA).

### Short tandem repeat analysis

Short tandem repeat analysis was performed at BEX CO., LTD. Using the genomic DNA extracted from iPSCs generated from patients with TTTS (AF-TTTS-iPSCs) and TTTS (AF-TTTS-iPSCs), 16 microsatelite markers were amplified by PCR with microsatellite-specific primers.

### Whole-exome sequencing

Approximately 2.0 μg of genomic DNA from each cell sample was sonicated to provide an average fragment size of 200–300 bp on a Covaris S220 instrument. After 5 cycles of PCR amplification, capture and library preparation were performed with Agilent SureSelect Human All Exon V5 + lincRNA (50 Mb), followed by washing, elution, and additional 12-cycle PCR amplification to attach index adaptors. Enriched libraries were sequenced on an Illumina HiSeq 2500 operated in 100-bp paired-end mode. Image analyses and base calling on all lanes of data were performed using bcl2fastq 1.8.4 with default parameters.

### Read mapping and variant analysis

Reads from the sample were first trimmed by removing adapters using cutadapt 1.7.1 and low-quality bases at ends using a custom script. Then, they were aligned to the hs37d5 sequence (hg19 and decoy sequences) using the Burrows–Wheeler Aligner 0.7.10. Mapped reads were converted from SAM to BAM using SAMtools 1.2 and processed by Picard 1.109 to eliminate PCR duplicate reads. The Genome Analysis Toolkit (GATK) 3.4 was then used to perform local realignment with known indel sites and map quality score recalibration to produce calibrated BAM files. Multi-sample callings for single-nucleotide variants (SNVs) were made by GATK. The annotated variant call format files were then filtered using GATK with a stringent filter setting and custom scripts. Annotations of detected variants were made using ANNOVAR based on GRCh37. Genotypes of control samples were shuffled for each variant from a perspective of protection of personal information.

### Teratoma formation

iPSCs were harvested by accutase treatment, collected into tubes, and centrifuged. The same volume of basement membrane matrix (BD Biosciences, cat. no. 354234) was added to the cell suspension. The cells (>1 × 10^7^) were subcutaneously inoculated into immunodeficient nude mice (BALB/cAJcl-*nu/nu*) (CREA, Tokyo, Japan). After 6–8 weeks, the resulting tumors were dissected and fixed with PBS containing 4% PFA. Paraffin-embedded tissue was sliced and stained with hematoxylin and eosin. The operation protocols were accepted by the Laboratory Animal Care and the Use Committee (A2003-002-C13-M05).

### Fluorescence-activated cell sorting analysis

The expression of cell surface markers on AFCs cultured in the amniomax II at passage 3 and keratinocytes derived from iPSCs were analyzed by flow cytometry. The AFCs and iPSC-derived keratinocytes were harvested by Trypsin-EDTA solution treatment and fixed with 2% PFA/PBS for 15 min at RT. After PBS wash, cells were permeabilized with 0.1% saponin (Wako, cat. no. 192-08851) and blocked with 5% goat serum. Primary antibodies were incubated for 1 h in PBS with 1% bovine serum albumin. After washing with PBS, cells were incubated with fluorescently coupled secondary antibodies; anti-rabbit or anti-mouse IgG conjugated with Alexa 488 (1:1000) for 30 min at room temperature. Secondary stain cells were analyzed on a BD LSR Fortessa (BD Biosciences).

### Protocol for differentiating iPSCs into keratinocytes

To induce differentiation, small clumps of undifferentiated iPSCs were subcultured onto a VTN-coated 10-mm dish in E8 medium on day 1 (protocol A). For comparison of differentiation protocols, single cells of iPSCs were subcultured onto a VTN-coated circle patterned CytoGraph (Dai Nippon printing Co., Ltd.) dish in E8 medium-supplemented 10-μM Y-27632 (Wako, cat. no. 251-00514) (protocol B) on day1. iPSCs were then incubated in defined keratinocyte serum-free medium (DKSFM) (Invitrogen, cat. no. 10744-019) supplemented with 1 μM all-trans retinoic acid (RA) (Wako, cat.no. 182-01111) and 10 ng/ml bone morphogenetic protein 4 (BMP4) (R&D systems. cat. no. 314-BP-010/CF) for 4 d. After 4 d, iPSCs were maintained in DKSFM supplemented with 20 ng/ml EGF (R&D systems, cat. no. 236-EG-200) until 14 days and passaged onto a 0.03 mg/ml coating of type I collagen (Advanced Biomatrix, cat. no. 5005-B) and 0.01 mg/ml fibronectin (Sigma Aldrich, cat. no. F0895-1MG)-coated dish in DKSFM supplemented with 10 μM Y-27632 (Wako, cat. no. 251-00514) and 20 ng/ml EGF. iPSC-derived keratinocytes were seeded at 3 × 10^4^/cm^2^ cells and enriched by rapid adherence to fibronectin and type I collagen-coated dishes for 15–30 min at room temperature. Nonadherent cells were discarded, and rapidly attached cells were cultured. EB method (protocol C) was performed as described previously (Bilousova et al., 2011). Embryonic bodies (EBs, n=100) were formed from 5 × 10^4^ iPSCs on a Petri dish in embryonic stem cell medium without basic fibroblast growth factor (ESC-no-bFGF medium). After 2 days, 30 EBs were transferred to a new Petri dish with ESC-no-bFGF medium containing 1-μM RA for three days in suspension culture. EBs were then transferred onto a type-IV (Sigma-Aldrich, cat. no. C7521-10MG)-coated 100-mm dish with ESC-no-bFGF medium containing 25-ngml^−1^ BMP4. After 3 days, the medium was switched to DKSFM, and the plated EBs were cultured for six days. EB remnants were removed by vacuum aspiration, and iPSC-derived keratinocytes were subcultured onto type-IV-coated 100-mm dish and selected rapid attachment to type-IV-coated dish for 15 minutes at room temperature in DKSFM.

### Generation of 3D skin equivalent

3D skin was generated according to a previously described protocol (Tsunenaga et al., 1994). To prepare the dermal equivalent, type I collagen (Koken Co., cat. no. IPC-50) and DMEM plus 10% FBS containing 1 × 10^6^ HFF2s were mixed and poured into an untreated 60-mm Petri dish while cooling and allowed to gel at 37°C for 1 h. Contraction of the collagen gel was facilitated by pulling the gel from the surface of the Petri dish. The medium was changed every 2 or 3 d for 7 d. HDK1-K4DT or iPSC-derived keratinocytes were plated at 2 × 10^5^ or 1 × 10^6^ cells inside in a glass ring (10 mm diameter) on the surface of the contracted collagen gel, which was plated onto polyethylene terephthalete membranes (Corning, cat. no. 35-3493). iPSC-derived keratinocytes were grown in DKSFM supplemented with 10 μM Y-27632 and 20 ng/ml EGF for 2 days, following which they were exposed to air in a 1:1 mixture of KSFM and DMEM plus 10% FBS, in which the Ca^2+^ concentration was adjusted to 1.8 mM. The medium was changed every 2 or 3 days. Multilayered 3D cultures of keratinocytes were obtained by days 14–21.

### Animal preparation and RA exposure

The procedure for creating MMC defects in fetal rats was based on the protocol described earlier (Danzer et al., 2005; Watanabe et al., 2010). Briefly, time-dated Sprague-Dawley rats (CLEA Japan, Tokyo or Sankyo Labo Service Co, Tokyo) were used. After being exposed to isoflurane (Wako, cat. no. 099-065-71), anesthetized rats were fed 60 mg/kg all-trans RA (Wako, Tokyo, cat. no. 182-01111) dissolved in olive oil (10 mg/ml) at embryonic day 10 (E10).

### Surgical procedure

Fetal intervention of RA-exposed Sprague-Dawley rats was performed at E20. Pregnant rats were anesthetized by isoflurane, and an abdominal midline incision was made to expose the uterine horns. The MMC defect was confirmed through the uterus under direct vision, and a small incision of the uterine wall and amniotic membranes was made directly above the defected site. 3D skin was transplanted into the area of defected skin. Beriplast P Combi-Set (CSL Behring, cat. no. 87799) was used for tissue adhesion. The hysterectomy site was closed by purse-string suture with 6-0 silk (Natsume Seisakusho, cat. no B10-60). The uterus was returned to the abdomen, and the abdominal incision site was closed by running suture. The fetuses were harvested by cesarean section at E22. Pregnant rats were then euthanized by cervical dislocation under anesthesia with isoflurane. The transplantation site was dissected and processed for histological and immunohistochemistry analysis. The primary antibody list is provided in Supplemental Table S2. Appropriate Alexia488 or alexa594-conjugated secondary antibody was used with DAPI nuclear counterstain. The operation protocols were accepted by the Laboratory Animal Care and the Use Committee (A2015-003-C01).

### Statistical analysis

Each experiment was repeated at least three times, and the data are presented as the mean ± SD of the mean. Statistical significance was determined by the student’s *t* test. *p* < 0.05 was considered statistically significant.

## Results

### Derivation and characterization of AFCs in patients with polyhydramnion

To examine whether iPSCs could be efficiently generated from human AFCs derived from patients with the serious disease coexisting polyhydramnion, we focused on two fetal diseases, namely TTTS and Down syndrome. Human AFCs were isolated from patients with TTTS (TTTS-AFC) and Down syndrome (T21-AFC) with the consents of subjects and the Ethical Review Board of the National Research Institute for Child Health and Development. AF was obtained through amniocentesis under sterile conditions during amnioreduction for therapy of polyhydramnion. Approximately 20 ml of AF is sufficient to obtain AFCs. The average number of viable cells was 0.356 × 10^6^/ml ± 0.227 × 10^6^/ml (mean ± standard deviation [SD]). The AFCs showed heterogeneous morphologies and reached confluence by 10 days after cell seeding (Figure 1A). Flow cytometry at passage 3 revealed that CD29, CD44, CD73, and human leukocyte antigen (HLA)-ABC [major histocompatibility complex (MHC) class I], were strongly positive; CD117 was rarely positive (0.8%), whereas CD14, CD19, CD34, HLA-DR, DP, and DQ (MHC class II) were negative (Figure 1B). Quantitative real-time polymerase chain reaction (qRT-PCR) analysis revealed trace amounts of OCT3/4, NANOG, and SOX2, compared with endometrial-derived mesenchymal stem cells (EDOM-MSCs) (Figure 1C). These results indicate that AFCs derived from polyhydramnion have a similar population to a normal volume of AF (De Coppi et al., 2007; Li et al., 2009; Tsai et al., 2004; You et al., 2008).

**Figure 1:**
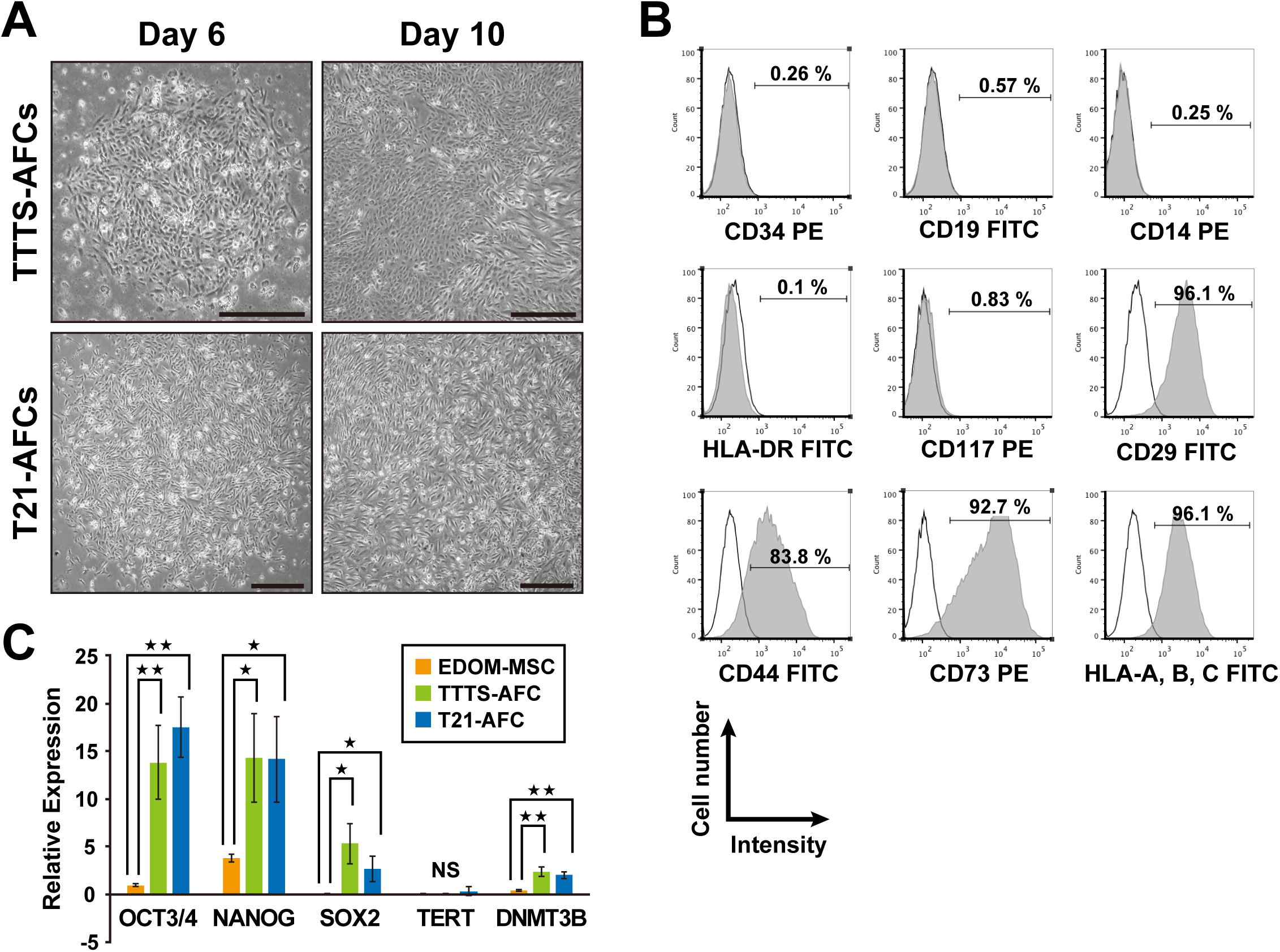
Characterization of human amniotic fluid cells (AFCs) derived from patients with Down syndrome and twin–twin transfusion syndrome (TTTS)-associated polyhydramnion. (A) Phase-contrast microscopic analysis for cell morphology in AFCs from patients with TTTS (TTTS-AFCs) and Down syndrome (T21-AFCs). Mesenchymal stem cell-like colonies appeared on the sixth day of culturing and 10^th^ day of culturing. Scale bar is 500 μm. (B) Flow cytometric analysis for CD29, CD44, CD73, CD117, CD14, CD19, CD34, human leukocyte antigen (HLA)-ABC, and HLA-DR. Isotype controls are shown in each panel. (C) Quantitative RT-PCR analysis for expression of OCT3/4’ NANOG, SOX2, TERT, and DNMT3B. Values are shown as mean ± standard deviation from three independent experiments.

### Generation of human iPSCs from AFCs of patients with TTTS and Down syndrome

AFCs at passages 3–4 were transfected with episomal vectors carrying six reprogramming factors (L-MYC, KLF4, OCT4, SOX2, LIN28, and short-hairpin RNA for P53) by electroporation. After electroporation, AFCs were subcultured into VTN-coated plates in chemically defined and serum-free Essential 8 medium. We observed the appearance of human embryonic stem cell (hESC)-like colonies 30–50 days after electroporation. Colonies with human ESC-like morphology expanded and grew as flat colonies with large nucleo-cytoplasmic ratios with a high level of alkaline phosphatase activity (Figure 2A). No significant differences in proliferation rates were detected between AF-T21-iPSCs and AF-TTTS-iPSCs. Neither cessation of cell proliferation such as senescence nor apoptosis/cell death was detected during cultivation through 70 passages. The reprogramming efficiency was 0.1% and 0.3% in AF-TTTS-iPSCs and AF-T21-iPSCs, respectively.

**Figure 2:**
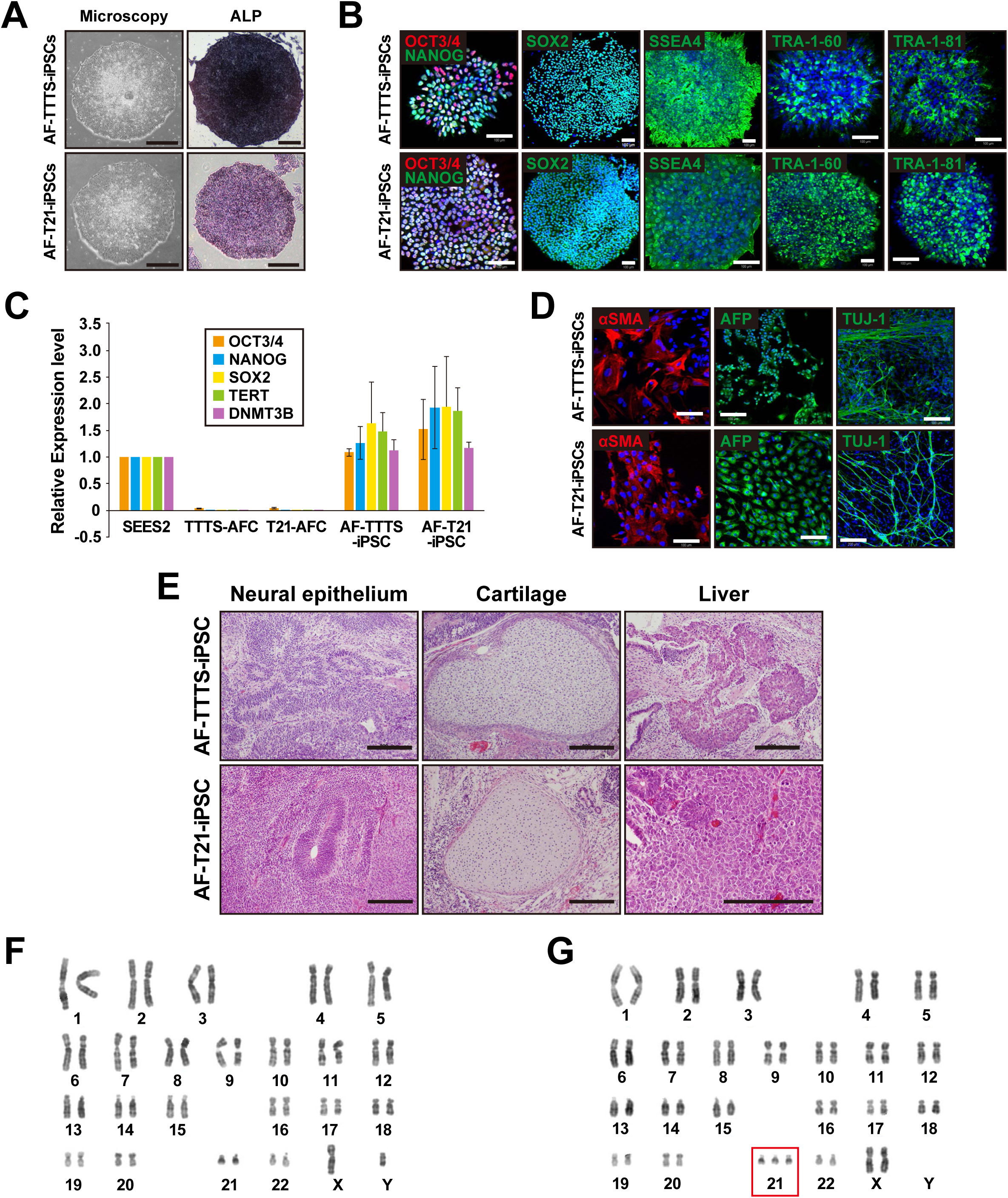
Generation of iPSCs from patients with Down syndrome (T21) and twin–twin transfusion syndrome (TTTS). (A) Phase contrast microscopic view of AF-TTTS-iPSC and AF-T21-iPSCs with embryonic stem cell (ESC)-like morphology growing on a feeder-free vitronectin surface (left panels). AF-TTTS-iPSCs and AF-T21-iPSCs are positive for alkaline phosphatase staining (ALP) (right panels). Scale bar is 500 μm. (B) Immunocytochemical analysis of stem cell markers, i.e. OCT3/4, NANOG, SOX2, SSEA-4, TRA-1-60, and TRA-1-81 in AF-TTTS-iPSCs and AF-T21-iPSCs colonies. AF-TTTS-iPSCs and AF-T21-iPSCs expressed these pluripotent markers. Nuclei were stained with DAPI (blue). Scale bar is 100 μm (C) qRT-PCR analysis on the endogenous expression levels of OCT3/4, NANOG, SOX2, TERT, and DNMT3B in AF-TTTS-iPSCs and AF-T21-iPSCs. The expression levels of these stem cell markers in AF-TTTS-iPSCs and AF-T21-iPSCs are comparable to those in human ESCs (SEES2). (D) *In vitro* differentiation of AF-TTTS-iPSCs and AF-T21-iPSCs into three germ layers. After embryoid body formation, iPSCs were stained with antibodies to α-smooth muscle actin (αSMA) (a mesodermal marker), α-fetoprotein (AFP) (an endodermal marker), and beta-III tubulin (TUJ-1) (an ectodermal marker). Scale bar is 100 μm. (E) *in vivo* differentiation of AF-TTTS-iPSCs and AF-T21-iPSCs into three germ layers. Teratomas were harvested 6–8 weeks after subcutaneous injection of iPS cells into nude mice. Various tissues, such as neural epithelium (ectodermal), cartilage (mesoderm), and liver (endoderm), were found. Scale bar is 200 μm. (F) Karyotypic analysis in AF-TTTS-iPSCs. AF-TTTS-iPSCs had normal karyotypes (46, XY). (G) Karyotypic analysis in AF-T21-iPSCs. AF-T21-iPSCs had typical trisomy karyotypes (47, XX, +21).

### Characterization of AF-T21-iPSCs and AF-TTTS-iPSCs

Both AF-TTTS-iPSCs and AF-T21-iPSCs expressed multiple pluripotency markers, including nuclear transcription factors OCT3/4, NANOG, SOX2, as well as surface antigen stage-specific embryonic antigen 4 (SSEA-4) and tumor-related antigen (TRA)-1-60 and TRA-1-81 (Figure 2B). qRT-PCR analysis showed that endogenous pluripotency marker genes, including OCT3/4, SOX2, NANOG, telomerase reverse transcriptase (TERT), and DNA methyltransferase 3 beta (DNMT3B), were activated in human iPSCs to a similar extent of those in control hESCs (SEES2) (Figure 2C), and the transgenes were fully silenced in AF-T21-iPSCs and AF-TTTS-iPSCs. We next examined the ability for *in vitro* differentiation by examining the expression of germ layer-specific markers in the embryonic body formation. Our analysis demonstrated that AF-T21-iPSCs and AF-TTTS-iPSCs were capable of differentiating into all three germ layers *in vitro* (Figure 2D). To examine pluripotency of iPSC clones, teratoma formation was performed by implantation of AF-T21-iPSCs and AF-TTTS-iPSCs at the subcutaneous tissue of immunodeficient nude mice. Three independent AF-T21-iPSCs and AF-TTTS-iPSCs clones induced teratomas within 6–8 weeks after implantation. Histological analysis of paraffin-embedded sections demonstrated that all three primary germ layers were generated as shown in Figure 2E. Thus, all AF-T21-iPSCs and AF-TTTS-iPSCs clones examined had potential for multilineage differentiation *in vivo.* Although teratoma derived from AF-T21-iPSCs showed the presence of neuroblastoma-like tissue (Supplemental Figure S1A) and liver tissue with a vacuolar structure (Supplemental Figure S1B), Down syndrome-specific findings were not detected. Karyotypic analyses revealed that the AF-TTTS-iPSCs clones had normal karyotypes (Figure 2F), whereas the AF-T21-iPSCs clones had trisomy 21 (Figure 2G). Analysis of 16 patterns of the short tandem repeat (STR) site revealed that all STR sites of AF-TTTS-iPSCs and AF-T21-iPSCs matched to those of their parental TTTS-AFC and T21-AFC (Supplemental Figure S1C).

### Whole-2 exome analysis of T21-iPSCs

The whole-exome analysis, wherein our sample was compared with the GRCh37 reference sequence, detected a heterozygous single-nucleotide variant (SNV) in the *CRELD1* focus. The C-to-G substitution (rs2302787), which results in a Pro-to-Arg alteration, was situated in exon 4. Several mutations of *CRELD1* have been reported to contribute to occurrence of cardiac atrioventricular septal defects in Down syndrome (Maslen et al., 2006). To know whether the alteration is deleterious, SIFT and Polyphen2 were employed. The former makes influence from similarity of amino acid sequences and gives scores close to zero when a variant is damaging, whereas the latter predicts effects of not only sequences but also 3D structures and provides scores close to 1.0 when a variant is intolerant. The scores for the variants were 0 and 0.999, respectively, suggesting a notable variant. Although its global allele frequency was 1.0%, higher frequency of 4.5% was documented for the Japanese population in the 1000 Genomes project (Supplemental Table S3).

### Generation and characteristics of iPSC-derived keratinocytes

We first attempted to generate iPSC-derived keratinocytes (iPSC-KC) based on the prior differentiation protocol (Bilousova et al., 2011; Guenou et al., 2009; Itoh et al., 2011; Metallo et al., 2008; Veraitch et al., 2013) using RA to promote ectodermal fate and BMP4 to block neural fate. To define the effective differentiation protocol, we compared differentiation efficiencies among three different protocols including direct differentiation using VTN-coated dish (protocol A), CytoGraph-coated dish (protocol B), and embryoid body method (protocol C) (Figure 3A). Protocols A and B were differed in coating agent. In protocol A, we modified the previously reported protocol (Itoh et al., 2011) by replacing matrigel with a human recombinant protein using VTN.

**Figure 3:**
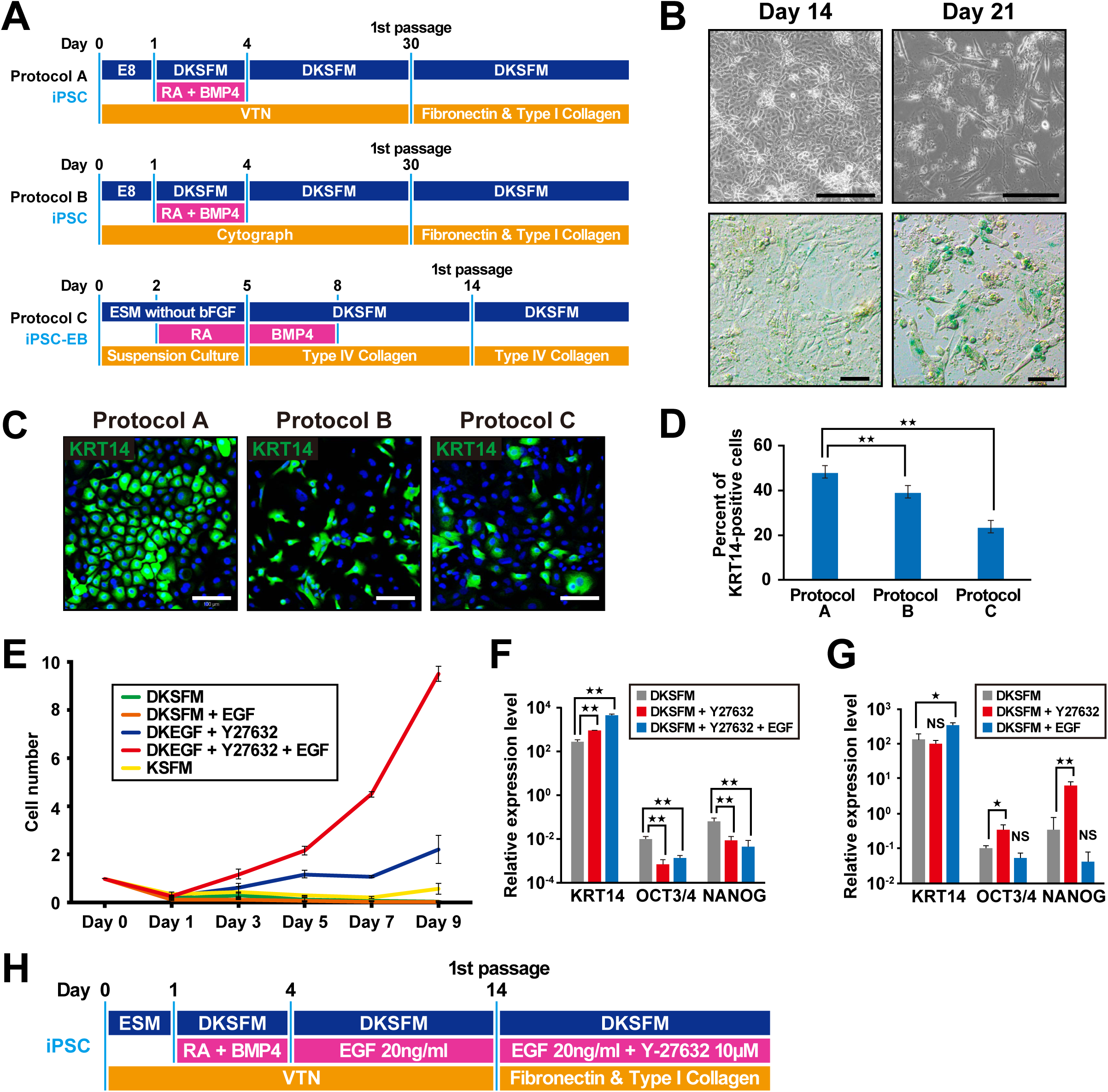
Establishment of differentiation protocol of iPSCs into the lineage of keratinocytes. (A) A schematic of the three differentiation protocols for generation of keratinocytes from iPSCs. Protocols A and B differed in the coating agents. Protocol C was performed via embryoid body (EB) formation (iPSC-EB). DKSFM, defined keratinocyte serum-free medium; RA, retinoic acid; BMP4, bone morphogenetic protein 4; VTN, vitronectin; E8, Essential 8; ESM, ESC medium. (B) Beta-galactosidase staining of iPSC-KCs at the indicated time points (days 14 and 21). Cell senescence was observed at day 21 of the induction. Scale bar is 100 μm. (C) iPSC-derived keratinocytes with different methods (protocols A, B, and C) at passage 2 were immunocytochemically stained with anti-keratin 14 (KRT14) antibody. Homogenous keratinocyte-like cells were stained in protocol A. Scale bar is 100 μm. (D) The number of cells positive for KRT14. The number of KRT 14-positive cells were highest in protocol A (48.08%), compared with protocol B (39.24%) and protocol C (23.62%). Data are shown as mean ± SD of the cell number from three independent experiments. KRT14: keratin 14. ^★★^*P* < 0.01. (E) The growth rate of iPSC-derived keratinocytes with different treatment. A combination of Y-27632- and epidermal growth factor accelerated cell growth. EGF, epidermal growth factor; DKSFM, defined keratinocyte serum free medium; KSFM, keratinocyte serum free medium. (F) qRT-PCR analysis of KRT14 and stem cell markers, i.e. OCT3/4 and NANOG, in iPSC-KCs at passage 2. KRT14 expression increased by the addition of epidermal growth factor (EGF). Data shown are mean ± SD of the expressions from three independent experiments. (G) qRT-PCR analysis of KRT14 and stem cell markers in iPSC-KCs at passage 0. Gene expression levels of OCT3/4 and NANOG increased in the presence of Y-27632, whereas the KRT14 expression level remained unchanged by the treatment of Y-27632. Data are presented as a mean ± SD of three independent experiments. (H) A schematic of the final protocol design for differentiation of iPSCs into keratinocytes. iPSC-KC, iPSC-derived keratinocytes; KSFM, keratinocyte serum-free medium; DKSFM, defined keratinocyte serum-free medium; RA, retinoic acid; BMP4, bone morphogenetic protein 4.

During direct differentiation (protocols A and B), cell senescence was observed at day 30, and these cells could not proliferate after the first passage. The number of keratinocyte-like cells decreased after 17 days and β-galactosidase staining revealed that cellular senescence was observed over 17 days, resulting in an exacerbated cellular state (Figure 3B). Therefore, the first passage was performed at 14–17 days in protocols A and B, respectively.

### Effect of Y-27632 on iPSC-KCs

Although we detected keratinocyte-like cells during passage 1 of protocols A, B, and C, we examined additional factors to obtain a sufficient number of iPSC-KC and found that Y-27632 was critical. Y-27632 is an inhibitor of Rho kinase that increases the long-term proliferative capacity of primary keratinocytes and promotes the differentiation of human bone marrow MSCs into keratinocyte-like cells (Chapman et al., 2010; Chapman et al., 2014; Li et al., 2015; Suprynowicz et al., 2012). Keratinocytes derived from iPSCs grown in the presence of Y27632 showed improved cell growth. The comparison of immunostaining of Keratin 14 (KRT14) at passage 2 with protocols A, B, and C revealed that the percentage of KRT14-positive cells reached 48.08%, 39.24%, and 23.62% in protocol A, B, and C, respectively, indicating that protocol A is most suitable for iPSC-KC proliferation (Figure 3C, 3D). Therefore, during subsequent experiments, differentiation of iPSCs into keratinocytes was performed by protocol A.

### Effect of EGF on iPSC-KCs

iPSC-KCs treated with Y-27632 showed improved cell growth; however, cell proliferation remained insufficient, required more than 2 weeks to obtain a sufficient cell number. To promote cell proliferation with a high expression level of epithelial markers, we examined iPSC-KCs to investigate the effect of EGF. iPSC-KCs treated with EGF and Y-27632 for 9 days after first passage (starting cell number was 1 × 10^5^/well) showed markedly improved cell growth compared with the cells treated with Y-27632, whereas the cell number of EGF-treated iPSC-KCs without treatment of Y-27632 did not proliferate at all, indicating that the combination of EGF and Y-27632 is important for the cell growth of iPSC-KCs (Figure 3E). Although iPSC-KCs grown in KSFM showed a higher cell growth than that grown in DKSFM (Figure 3E), we used DKSFM for subsequent experiments as it is chemically defined and optimized for growth and expansion of human keratinocytes without the potential contamination derived from animal serum.

We further analyzed gene expression levels of KRT14 in iPSC-KC at passage 1 and found that iPSC-KCs treated with EGF and Y-27632 expressed KRT14 at a higher level than those treated with Y-27632 alone (Figure 3F). Flow cytometric analysis also showed that KRT14-positive population increased in cell number and intensity by addition of Y-27632 and EGF (Supplemental Figure S2A). Treatment with EGF for day 4 to 14 resulted in a marginal yet significantly higher expression of KRT14 (Figure 3G). Y-27632 was not included for day 4 to 14 because Y-27632-treated iPSC-KCs before the first passage showed a higher expression of OCT3/4 and NANOG (Figure 3G). SSEA4-positive undifferentiated iPSCs were successfully removed by treatment of EGF and Y-27632 after the first passage as confirmed by flow cytometric analysis (Supplemental Figure S2B). As for the growth medium, DKSFM was used throughout the differentiation process (Supplemental Figure S2C). Taken together, we concluded that differentiation efficiency of iPSC-KCs was most effective in protocol A using DKSFM supplemented EGF and Y-27632, as shown in Figure 3H.

### Characterization of keratinocytes derived from iPSCs

In our protocol, overall induction periods were further reduced compared with previously reported protocols (Guenou et al., 2009; Itoh et al., 2011; Itoh et al., 2013), and iPSC-KCs were successfully increased and could be expanded for greater than five passages at least. Interestingly, in terms of proliferative and differentiation abilities, both Y-27632 and EGF were required for the generation of iPSC-KC. Y-27632 was more of affecting the proliferative ability, and EGF was more of affecting the terminal differentiation. Keratinocytes derived from AF-TTTS-iPSCs (TTTS-iPSC-KCs) and AF-T21-iPSCs (T21-iPSC-KCs) showed keratinocyte-like morphology at passage 4 like HDK1-K4DT (Figure 4A). KRT14 gradually increased for each passage (Figure 4B). Transcriptional analysis by qRT-PCR showed that the gene expression levels of KRT14, KRT18, p63, OCT3/4, and NANOG of TTTS-iPSC-KCs and T21-iPSC-KCs at passage 4 reached to those in HDK1-K4DT (Figure 4C). Interestingly, T21-iPSC-KCs showed higher expression of terminal differentiation markers such as INVOLUCRIN and FILAGGRIN than TTTS-iPSC-KCs (Figure 4D). This result is consistent with the clinical features of hyperkeratosis that are frequently observed in patients with Down syndrome. Immunostaining revealed the expression of KRT14, p63, and laminin 5 both in TTTS-iPSC-KCs and in T21-iPSC-KCs (Figure 4E). Moreover, keratin 10 (KRT10) and involucrin, markers of differentiated keratinocytes of the suprabasal layer, were induced under a high calcium condition. As expected, the number of involucrin-positive T21-iPSC-KCs increased under a low calcium condition. Keratin 15 (KRT15), a marker of epithelial stem cells in the hair follicle, was rarely detected (Figure 4E). The KRT14-positive cell population in EGF and Y-27632-treated iPSC-KC at passage 2 reached to 95.9%, as shown by flow cytometric analysis (Figure 4F).

**Figure 4:**
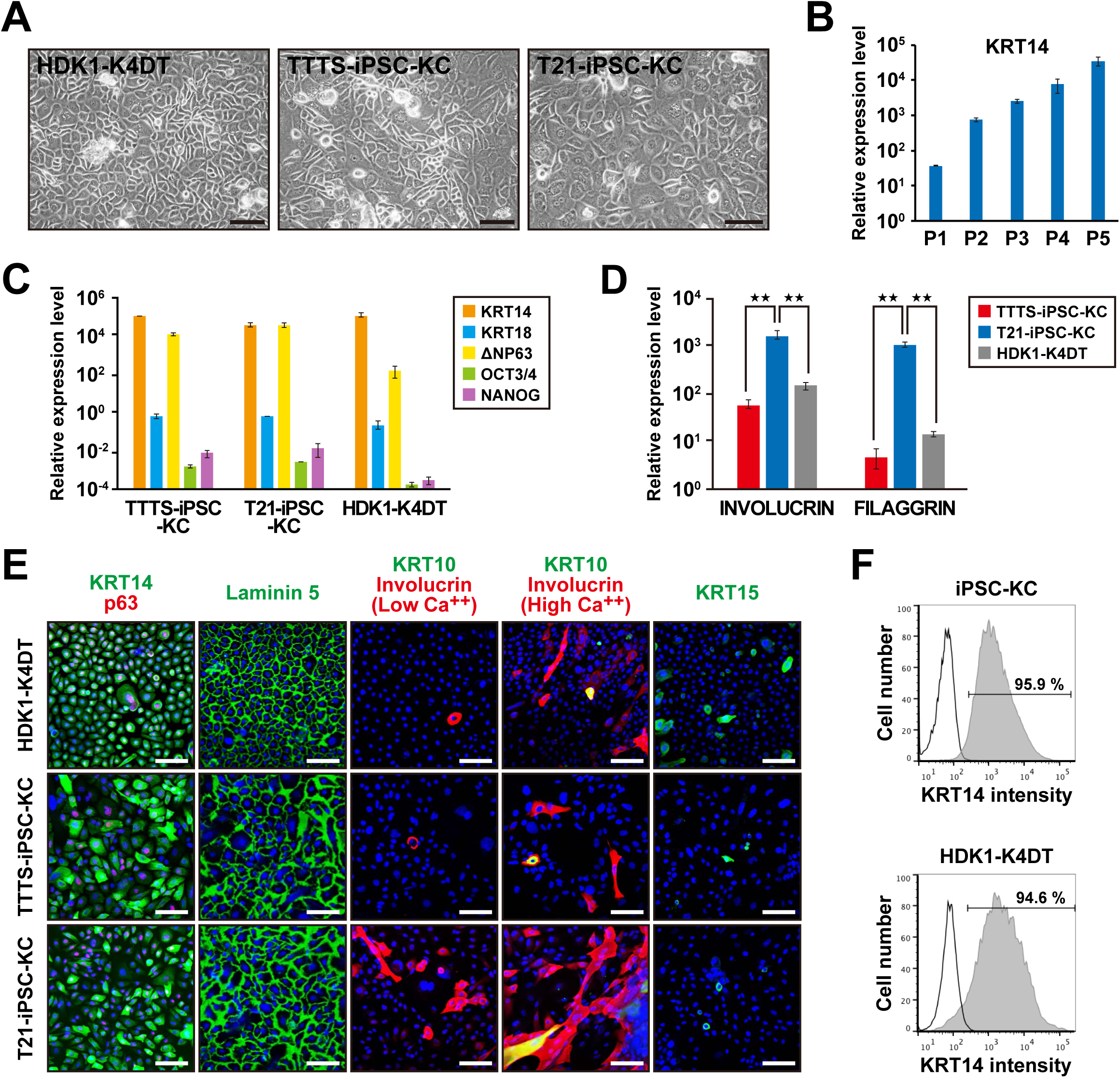
Characterization of a homogeneous population of keratinocytes derived from induced pluripotent stem cells (iPSCs). (A) Phase-contrast microscopic analysis of human dermal keratinocytes (HDK1-K4DT) and keratinocytes derived from AF-T21-iPSCs (T21-iPSC-KC) and AF-TTTS-iPSCs (TTTS-iPSC-KC). Both iPSC-derived keratinocytes showed human keratinocyte-like morphology. Scale bar is 500 μm. (B) qRT-PCR analysis of keratin 14 (KRT14) at each passage. (C) qRT-PCR analysis of epithelial markers, i.e., KRT14, KRT18, and ΔNP63, and stem cell markers, i.e. OCT3/4 and NANOG, in human iPSC-KCs at passage 4. The epithelial marker expression levels in iPSC-KCs were similar to those of HDK1-K4DT. (D) qRT-PCR analysis of terminal differentiation markers, i.e. INVOLUCRIN and FILAGGRIN. T21-iPSC-KCs showed higher expression levels of INVOLUCRIN and FILAGGRIN than TTTS-iPSC-KCs did. (E) Immunofluorescence of epithelial markers, i.e. KRT14, p63, laminin 5, involucrin, KRT10, and KRT15. Under the high-calcium condition, iPSC-KCs expressed involucrin and KRT10 at a higher frequency, indicating accelerated epidermal differentiation. Scale bar is 100 μm. (F) Flow cytometric analysis of KRT14 in iPSC-KCs at passage 2. Isotype controls are included in each panel.

### Passage-dependent epidermal differentiation of iPSC-KCs for 3D skin

HDK1-K4DT cells had a strong proliferative ability with high expression levels of epithelial markers during the long-term culture period. iPSC-KCs reduced proliferative ability as each passage progressed; however, their terminal differentiation ability increased as each passage progressed (Figure 4B and Supplemental Figure S3A). Cell morphology of iPSC-KCs appeared spindle-like during early passages and became similar to human keratinocytes as the passages progressed (Supplemental Figure S3B). However, a vacuolar degeneration-like structure was observed at passage 5, and the cells ceased to proliferate (Supplemental Figure S3B).

For the successful construction of 3D skin with iPSC-KCs, we investigated proper passage number of iPSC-KCs with both sufficient expression levels of KRT14 and the ability of cells to grow. Unfortunately, 3D skin was not generated with iPSC-KCs by the same method as HDK1-K4DT. Thus, we increased the initial seeding cell number of iPSC-KCs from 2 × 10^5^ to 1 × 10^6^ and allowed 2 d for the culture period in iPSC-KCs before high Ca^2+^ induction at the air–liquid interface. iPSC-KCs at passage 0 constructed epidermis that were negative for ki67, KRT14, p63 and Pan-cytokeratin (Pan-CK) (Supplemental Figure S4). The 3D skin with iPSC-KCs at passage 1 revealed immature epithelial-like tissue that expressed pan-CK and Ki67 but not other epithelial markers, including KRT14 and P63 (Supplemental Figure S4). The 3D skin with iPSC-KCs at later passages (more than passage 4) showed mature keratinocytes with the prickle cell layer that were strongly positive for KRT14, Pan-CK, and involucrin, and weakly positive for Ki67 (Supplemental Figure S4). Finally, we successfully identified the most suitable condition to generate a multilayered epidermis with iPSC-KCs at passage 3. The 3D skin with iPSC-KCs at passage 3 after 14 d of air-liquid cultivation had a multilayered epidermis with the KRT14 in the basal compartment and laminin 5 at the dermal-epidermal junction (Figure 5). Moreover, KRT10 was detected in the suprabasal layer, and loricrin and filaggrin, late markers of epidermal differentiation, were also observed in the upper layers of the epidermis. These data suggest that AF-TTTS-iPSCs and AF-T21-iPSCs can be differentiated into fully functional keratinocytes for artificial 3D skin. This success may be attributed to the favorable balance of proliferation and differentiation capacities in iPSC-KCs.

**Figure 5:**
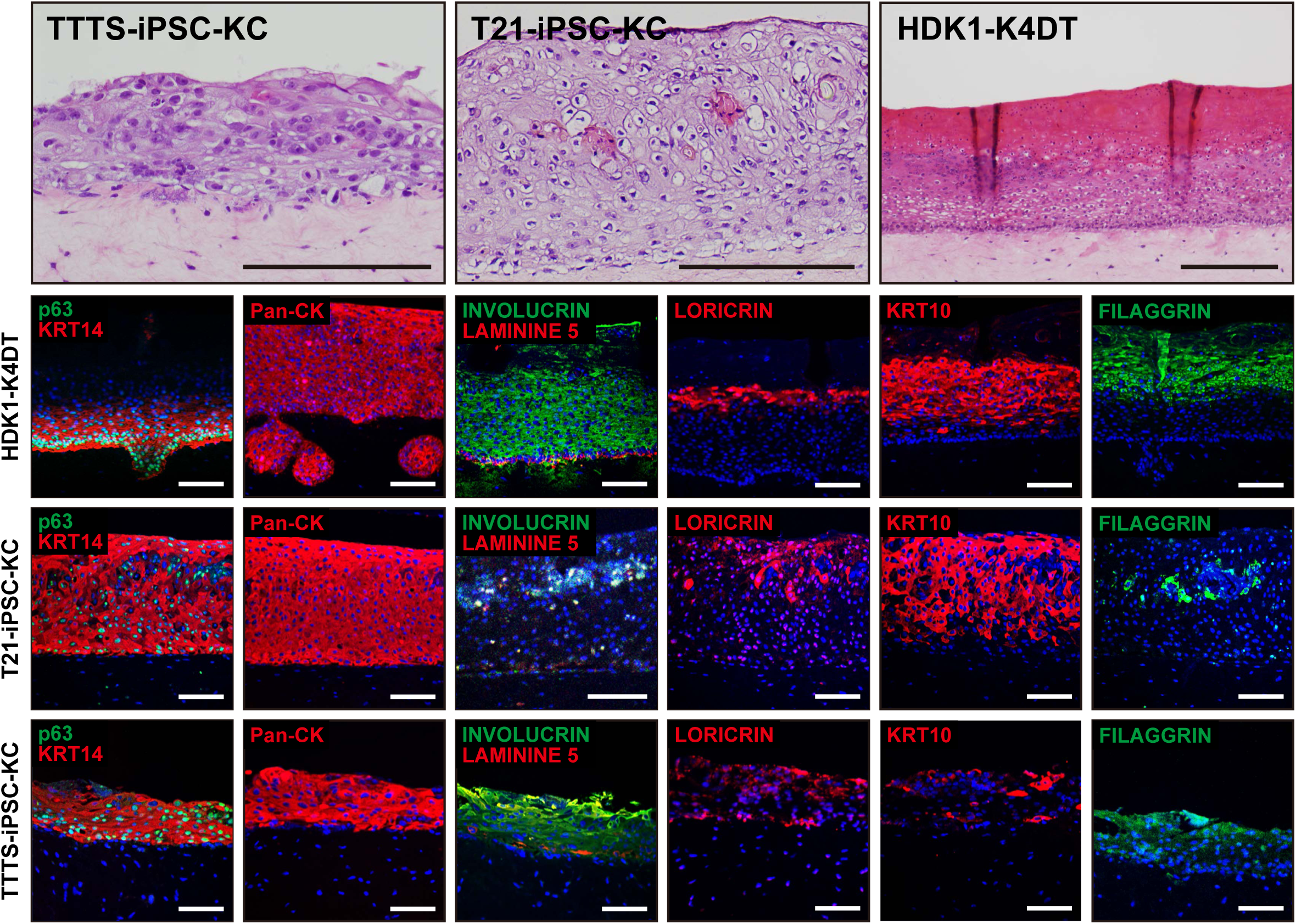
Three-dimensional (3D) cultured skin equivalents using iPSC-KCs. Upper panels: Histology of 3D skin with T21-iPSC-KCs, TTTS-iPSC-KCs, or HDK1-K4DT. Lower panes: Immunohistochemical analysis with antibodies to KRT14, p63, Pan-CK, involucrin, laminin 5, loricrin, KRT10, and filaggrin. The multilayered epidermis expressed KRT14, involucrin, laminin 5, Pan-CK, loricrin, KRT10, and filaggrin in artificial skin, indicating that iPSC-KCs terminally differentiate in these skin equivalents. Scale bar is 100 μm.

### Transplantation of 3D skin into MMC fetal rats

A total of 97 fetuses were viable after cesarean section. MMC was present in 83.5% (81 of 97) of fetuses exposed to RA. These MMC rats showed a defect in the skin and spine and exposure of the spinal cord (Figure 6A). Cross-sectional analysis confirmed the MMC defects and showed the degenerated spinal cord in rats exposed to RA (Figure 6B). In total, 20 fetal rats were operated on at day 20 of gestation. The 3D skin with iPSC-KCs was transplanted into fetuses across a small incision of the uterine wall following closure of the incision site (Figure 6C). A cesarean section was performed at day 22 of gestation. The survival of rats was lower in the fetal therapy group (15 of 20; 80%) compared with non-treated MMC fetus group (61 of 61; 100%). Twelve neonatal rats had a complete or partial skin defect coverage with 3D skin [complete coverage: 4 of 20 (20%), incomplete coverage: 8 of 20 (40%)], virtually isolating the spinal cord from direct exposure to the amniotic cavity (Figure 6D). Birth weight and crown-rump length (CRL) were significantly lower in the fetal therapy group than in normal rats and non-treated MMC rats (Figure 6E). Although the transverse length (TL) and vertical length (VL) of the MMC defect size were slightly shorter in the fetal therapy group than in fetuses without therapy, there were no significant differences when the TL and VL were adjusted for overall fetal CRL (Figure 6F).

**Figure 6:**
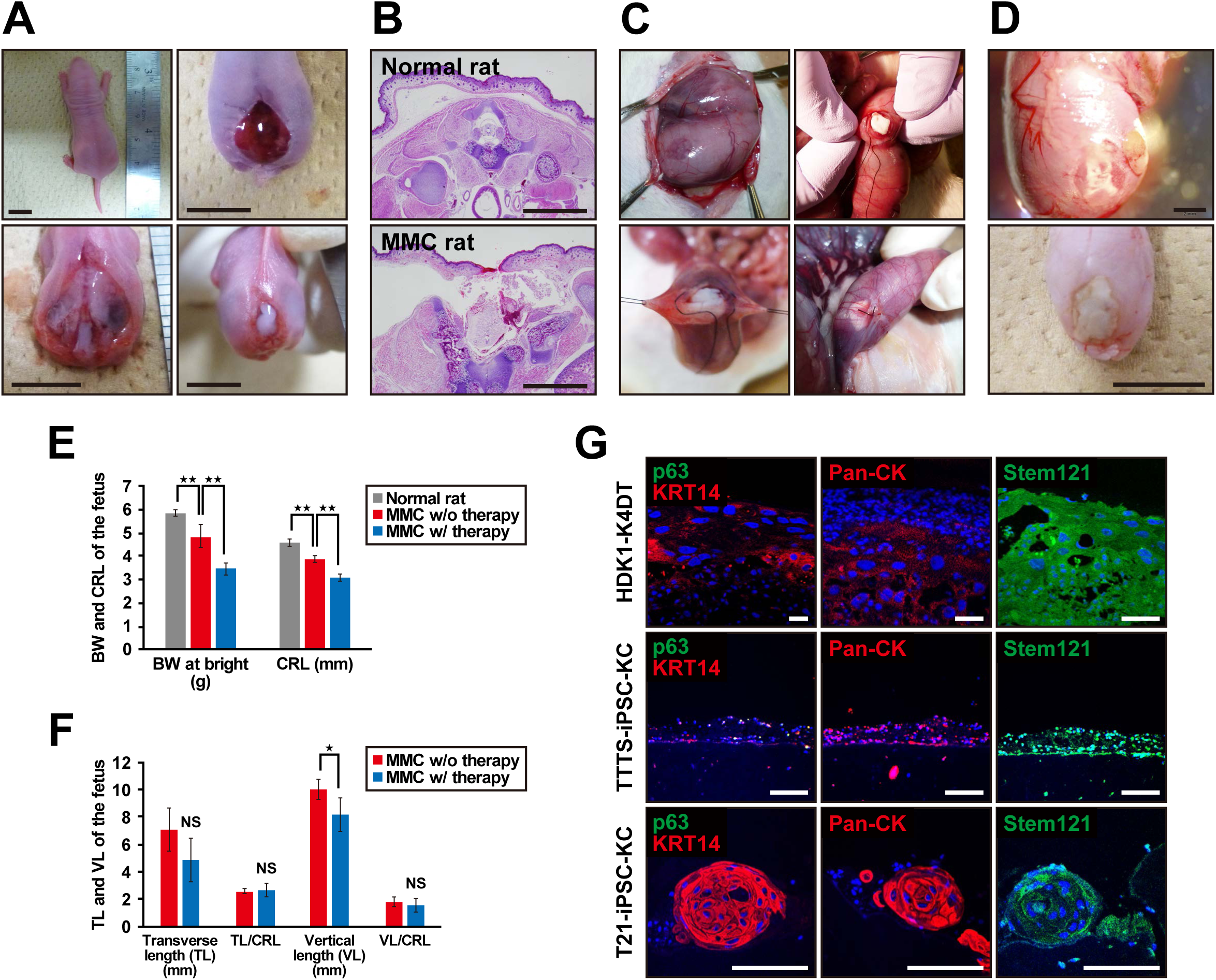
The rat model of myelomeningocele (MMC) with application of artificial skin. (A) Gross pathology images of the MMC lesion site at birth in a normal rat (upper-left panel) and MMC rat (upper-right and lower panels). MMC rat shows defects in skin (upper-right) and spine (lower-left) and exposure of spinal cord (lower-right). (B) Cross-sectional images of spinal cord at lumbar levels of a normal rat and MMC rat. Scale bar is 2 mm. (C) Gross view of the intrauterine transplantation of artificial skin. Representative images of major steps of the MMC repair process in the fetal rat model. A MMC defect site was identified through the uterus (upper-left panel). A small incision of the uterus was made just above the defect site, following which artificial skin was transplanted (upper-right panel). Finally, the uterine wall was closed over the defect (lower panels). (D) Gross views of neonatal rat with myelomeningocele (MMC) at birth (E22). Representative photograph in incomplete closure (upper panel) and complete closure (lower panel) after artificial skin application at day 20. (E) Birth weight (BW) and cranio-caudal length (CRL) of normal neonatal and neonatal MMC rats with or without fetal therapy. MMC rats with fetal therapy group were born with smaller BW and CRL compared to normal rats or non-treated MMC rats. ^★★^*P* < 0.01. (F) Comparison of MMC defect size between neonatal MMC rats with fetal treatment and rats without therapy. There were no significant differences in the MMC size by fetal therapy when adjusted for the CRL. TL, transverse length; VL, vertical length. ^★^*P* < 0.05. NS, not significant. (G) Expression profile of epidermal markers and short-term *in vivo* effect on regeneration of rat skin defect in artificial skin. Transverse section through the myelomeningocele defect 2 d after a transplantation of artificial skin with iPSC-KC epidermis. The expressions of epidermal markers, i.e. KRT14, p63, pan-cytokeratin (Pan-CK) and stem121, were analyzed in artificial skin with HDK1-K4DT, TTTS-iPSC-KCs, or T21-iPSC-KCs. Scale bar is 100 μm.

### Histological evaluation of 3D skin at transplantation site

Immunohistochemistry of the transplanted artificial skin with iPSC-KCs and HDK1-K4DT demonstrated that the expression of epithelial markers, such as KRT14 and p63 was weak, but obviously remained after birth (Figure 6G). Transplanted 3D skin was confirmed using human-specific antigen cytoplasmic protein detected with “stem121”. Moreover, epidermal ingrowth was detected at the edge of the epidermis defect in MMC rats, and the epidermis appeared to be elongated underneath the transplantation site (Supplemental Figure S5).

## Discussion

This study is designed to obtain preclinical proof of concept for fetal therapy to patients with MMC using autologous iPSCs. Pregnant women with polyhydramnion need to receive amnioreduction therapy, i.e. 1,000 to 2,000 ml of AF is usually aspirated for preventing threatened premature delivery and respiratory discomfort. A large number of AFCs can be obtained from 20 ml of AF by amnioreduction. The sufficient number of AFCs with their prominent proliferation capability leads to successful iPSC generation from patients with lethal disorders, such as TTTS.

### Artificial Skin with iPSC-epidermal cells and dermal fibroblasts

Epidermal sheet products are applied to patients with skin disorders, such as severe burn, giant congenital melanocytic nevus, vitiligo vulgaris, and epidermolysis bulla (Kamao et al., 2014; Kishi et al., 2010; Lu et al., 2014) and are often used with a combination with cryopreserved allogeneic skin, artificial dermis, non-surgical granulation tissue, meshed/patched autologous skin, and fresh allogeneic skin (Matsumura et al., 2016). Artificial skin with adequate strength is required because not only epidermis but also dermis is absent in the defect site of MMC. Therefore, epidermis alone is not sufficient to cover the defect.

In addition to the tissue engineering approach with usage of bioengineered products, organoid is formed in vitro in 3D as a miniaturized and simplified version of an organ. The organoid self-organizes in 3D culture through self-renewal and differentiation capacities of pluripotent stem cells, such as iPSCs and ESCs. Organoids, such as eye, gut, liver, and brain, are being developed (Hasegawa et al., 2016; Sasai, 2013; Takebe et al., 2012; Uchida et al., 2016; Yokobori et al., 2016). A skin organoid, however, has not been developed so far to assure strength to cover the defect site of MMC.

### Challenge of an iPSC therapy to diseased fetus

The first autologous cellular therapy was performed to a patient with macular degeneration with iPSCs generated from skin biopsy (Kamao et al., 2014). Herein, we hypothesize that AFC-derived iPSCs can be used for diseased fetus. The differentiation protocol developed in this study enables earlier induction of keratinocytes from iPSCs than previously reported. However, the window period of acceptable fetal therapy is usually 19–26 weeks of gestation due to fetal condition. Therefore, to perform cellular therapy during pregnancy, allogeneic products with appropriately selected haplotypes may be more feasible because even less than 100 cell lines of iPSCs from banks would be sufficient to cover 90% of the Japanese population (Nakatsuji et al., 2008).

### Efficient epidermal differentiation of iPSCs with rock inhibitor

The efficient keratinocytic differentiation in a short period was achieved by supplementation with both EGF and a rock inhibitor(Guenou et al., 2009; Itoh et al., 2011; Itoh et al., 2013). Y27632, a rock inhibitor, affects the differentiation and proliferation of various types of stem cells (Joo et al., 2012; Kurosawa, 2012; Watanabe et al., 2007). Moreover, Y27632 enables primary human keratinocytes not only efficiently to bypass senescence but also to increase long-term proliferation (Chapman et al., 2010; Chapman et al., 2014). Furthermore, Y27632 facilitates differentiation of mesenchymal stem cells into keratinocyte-like cells (Li et al., 2015). Likewise, Y-27632-treated iPSC-KCs showed improved cell growth and induced differentiation of human iPSCs into functional keratinocytes that were used for manufacturing of 3D skin.

EGF is the critical factor for the generation of human KRT15-positive epithelial stem cells, which are capable of differentiating into multiple cell types, such as cells of hair follicles and interfollicular epidermis and KRT14-positive basal cells from human iPSCs (Yang et al., 2014). These are compatible with the results in this study that showed an increased number of KRT14-positive iPSC-KCs and up-regulated expression of the KRT14 gene along with decreased expression of pluripotency markers. In contrast, lack of KRT15 in iPSC-KCs indicates that human iPSC-KCs generated with our protocol comprised a larger proportion of mature keratinocytes than KRT15-positive epithelial stem cells, which may decrease during the induction process. Along with the benefit of EGF, we have to be careful because our differentiation protocol using EGF may induce other types of skin stem cells under certain conditions in 3D culture due to induction of sebocytic lineage with EGF (Zouboulis, 2013).

### fetal therapy to a patient with MMC with 3D skin

Fetal therapy started with patients with TTTS, fetal anemia, and MMC (Adzick and Harrison, 1994; De Lia et al., 1990; Rodeck et al., 1981); however, a standard procedure has not yet been developed (Danzer and Johnson, 2014). As for MMC, laparoscopic surgery to suture skin defects is one of the fetal therapies so far in cases wherein skin defects are not large. Artificial skin needs to be developed due to coverage of large skin defects along with surgical approach. For generation of artificial 3D skin, iPSCs can be efficiently generated from AFCs derived from polyhydramnion in integration-free, xeno-free, and serum-free conditions because AFCs derived from patients with polyhydramnion exhibited prominent proliferative capacity *in vitro.*

An important limitation of this rat model is the short gestation period of rats without the ability to analyze the longer-term effects of artificial skin *in vivo.* Further studies to analyze the longer-term effect, including regeneration of skin, tumorigenic potential, and neurological prognosis, are required in large animal model. In the fetal therapy strategy, improvement in distal neurological function remains limited because of the failure to reverse neurological injury that occurred prior to the time of repair (Heffez et al., 1990). The transplantation of neural stem cells was reported to improve neurological outcome in an animal model of nerve injury, such as spinal cord injury (Biernaskie et al., 2007; Hu et al., 2010; Sieber-Blum, 2010; Wang et al., 2011). iPSC-derived neural crest stem cells integrate into the injured spinal cord in the fetal lamb model of MMC (Saadai et al., 2013). Additional cell types, such as neural stem cells/neural crest cells, to the artificial skin may be beneficial to a future fetal therapy.

In this study, we demonstrated a novel fetal MMC therapy that can be achieved by cellular therapy using AFC-derived iPSCs. Our fetal cell treatment is minimally invasive and therefore has the potential to become a novel treatment for MMC.

## Acknowledgments

This study was supported by a grant from JSPS KAKENHI (grant number JP15K19665). We are grateful to Dr. Tohru Kiyono, Division of virology chief at National cancer center research institute, for donating the cell line of HDK1-K4DT. We sincerely thank Minoru Ichinose (NCDHD) for performing sectioning. The encyclopedic pathological knowledge of Michoyo Nasu (NCDHD) was gratefully appreciated. I am also grateful to the Centre for Maternal-Fetal, Neonatal and Reproductive Medicine (NCCHD) for AF collection. The authors would like to thank Enago (www.enago.jp) for the English language review.

## Conflict of Interest

There are no conflicts of interest to declare.

## Supplementary Information

**Supplemental Figure S1. Teratoma formation of in amniotic fluid cells from patients with Down syndrome (AF-T21-iPSC) in vivo**

(A) Neuroblastoma-like tissue

(B) Liver tissue with vacuolar structure

(C) STR analysis of AF-T21-iPSC, T21-AFC, AF-TTTS-iPSC, and TTTS-AFC. Short tandem repeat (STR) profiling was performed by BEX CO., LTD, Tokyo, Japan. The 16 loci analyzed by the PowerPlex 1.2 system (Promega, Madison, WI, USA) was composed of D3S1358, TH01, D21S11, D18S51, Penta E, D5S818, D13S317, D7S820, D16S539, CSF1PO, Penta D, AMEL, vWA, D8S1179, TPOX, and FGA. T21-AFC and TTTS-AFC were parental cells of AF-T21-iPSC and AF-TTTS-iPSC, respectively.

**Supplemental Figure S2. Establishment of differentiation protocol of iPSCs into the lineage of keratinocytes**

(A) Flow cytometric analysis of KRT14 in iPSC-KCs at passage 2 (gray) and at passage 0 (open). iPSC-KCs were exposed to Y-27632 and EGF.

(B) Flow cytometric analysis of SSEA4 in T21-iPSC-KCs. T21-iPSC-KCs at passage 1 were exposed to Y27632 alone or Y27632 and EGF for one week and then applied to the flow cytometric analysis. Undifferentiated AF-T21-iPSCs were also shown in black for reference. Isotype control is included at each panel.

(C) qRT-PCR analysis of the KRT14 transcripts in iPSC-KCs maintained either in a medium of DKSFM or CNT-PR with EGF and Y27632. The KRT14 gene expression level was higher with the use of DKSFM than with the use of CNT-PR medium. Data are presented as a mean ± SD of three independent experiments.

**Supplemental Figure S3: Characterization of T21-iPSC-KCs at different passage number**

(A) The growth rate of T21-iPSC-KCs with the different passage number. Growth rate was reduced as each passage progressed. The initial cell number was 1 × 10^5^/well, and cell number was counted at the indicated days after cell seeding.

(B) Cell morphology of T21-iPSC-KCs in different passage number. iPSC-KCs at early passage exhibited spindle cell morphology (iPSC-KC at P1-2) and became keratinocytic (iPSC-KC at P3-4). In later passages, keratinocyte-like cells with a vacuolar degeneration (iPSC-KC at P5-6) were observed.

**Supplemental Figure S4. Immunohistochemical analysis on three-dimensional (3D) culture of iPSC-KCs at different passages**

iPSC-KCs at passage 0 were negative for KRT14, p63, pan-cytokeratin (Pan-CK), and Ki67. iPSC-KCs at passage 1 became positive for pan-CK and Ki67 but not for KRT14 and p63. iPSC-KCs at passage 5 were positive for KRT14, Pan-CK, and involucrin but not for p63, indicating terminal differentiation of epidermis in the 3D-culture.

**Supplemental Figure S5. Short-term in vivo effect on regeneration of rat skin defect with iPSC-KC artificial skin**

Epidermal ingrowth from the edge of MMC defect was observed underneath the artificial skin under low magnification (A) and high magnification (B). Scale bar is 500 μm.

